# Functional coupling between MC4R and Kir7.1 contributes to clozapine-induced hyperphagia

**DOI:** 10.1101/2024.06.07.597973

**Authors:** Li Li, Ciria C. Hernandez, Luis E. Gimenez, Baijie Xu, Naima S. Dahir, Swati, Shari G. Birnbaum, Roger D. Cone, Chen Liu

## Abstract

Most antipsychotic drugs (APDs) induce hyperphagia and weight gain. However, the neural mechanisms are poorly understood, partly due to challenges replicating their metabolic effects in rodents. Here, we report a new mouse model that recapitulates overeating induced by clozapine, a widely prescribed APD. Our study shows that clozapine boosts food intake by inhibiting melanocortin 4 receptor (MC4R) expressing neurons in the paraventricular nucleus of the hypothalamus. Interestingly, neither clozapine nor risperidone, another commonly used APD, affects receptor-ligand binding or the canonical Gαs signaling of MC4R. Instead, they inhibit neuronal activity by enhancing the coupling between MC4R and Kir7.1, leading to the open state of the inwardly rectifying potassium channel. Deletion of *Kir7.1* in *Mc4r-Cre* neurons prevents clozapine-induced weight gain, while treatment with a selective Kir7.1 blocker mitigates overeating in clozapine-fed mice. Our findings unveil a molecular pathway underlying the effect of APDs on feeding behavior and suggest its potential as a therapeutic target.

## INTRODUCTION

Antipsychotic drugs (APDs) demonstrate broad efficacy in treating various neuropsychiatric conditions ^1^. However, most of them cause drug-induced hyperphagia, leading to rapid weight gain ^2^. Despite efforts to implement lifestyle changes, nutritional consulting, and weight-loss medications ^3–5^, the metabolic side effects of APDs are often intolerable for many patients, becoming a leading cause of drug discontinuation.

APDs bind receptors for multiple neurotransmitters in the brain ^6^. While the psychotropic effects are mainly attributed to the blockade of dopamine D2 and serotonin 2a receptors ^7^, the neural mechanisms that underlie their adverse impact on food intake remain less clear. Genome-wide association and candidate gene studies in humans have implicated the melanocortin 4 receptor (*MC4R*) ^8^, among others ^9–12^, in APD-induced weight gain. *Mc4r* encodes the receptor for two hypothalamic neuropeptides that reciprocally regulate food intake: α-melanocyte-stimulating hormone (α-MSH)—a proteolytic product of pro-opiomelanocortin (POMC)—activates MC4Rs to suppress food intake, whereas the orexigenic agouti-related peptide (AgRP) also binds to the MC4Rs to stimulate food consumption ^13^. Of note, AgRP action is complex, acting as a competitive antagonist of α-MSH ^14,15^, an inverse agonist of the MC4R ^16^, and a biased agonist, coupling the MC4R to the opening of the ion channel Kir7.1 ^17,18^. Recent studies have shown that olanzapine, an APD, boosts food intake by up-regulating *Agrp* transcription in the hypothalamus. Additionally, olanzapine-induced weight gain was attenuated in mice lacking *Agrp* ^19^.

POMC and AgRP-expressing neurons integrate circulating metabolic cues and synapse on neurons in the paraventricular nucleus of the hypothalamus (PVH) that express *Mc4r* (designated hereafter as MC4R^PVH^ neurons). These neurons can bidirectionally regulate feeding: their activation reduces appetite, whereas inhibition of these neurons drives hyperphagia and weight gain ^20,21^. Interestingly, when applied to hypothalamic brain slices, risperidone, another APD, hyperpolarizes MC4R^PVH^ neurons via the opening of postsynaptic potassium channels ^22^. Furthermore, pretreatment with an MC4R antagonist prevented this inhibition. These findings suggest that risperidone interacts with MC4Rs and support the observation that *Mc4r* in PVH neurons is necessary for the drug’s effect on food intake and body weight ^22^.

Nevertheless, the specific potassium conductance that mediates risperidone-induced inhibition is unknown. Furthermore, whether other APDs act on the same pathway to induce hyperphagia remains to be determined. Notably, one particular challenge in studying APDs’ effect is the difficulty of recapitulating human metabolic syndrome in mice. For example, clozapine, one of the most effective APDs, causes excessive weight gain in both youths and adults ^23^. Yet, previous attempts to chronically administer clozapine to rodents via intraperitoneal injections or drinking water only slightly impact food intake and body weight ^24,25^. In the current study, we developed a new mouse model of clozapine-induced weight gain and studied the neural mechanisms behind the drug-induced hyperphagia.

## RESULTS

### Clozapine treatment causes excessive weight gain in female C57BL/6 mice

We formulated a clozapine diet by compounding clozapine (50 milligrams per kilogram) into a control diet (Research Diets, Inc. D09092903). The clozapine and control diets have the same macronutrient composition and energy density (**Table S1**). Additionally, mice exhibited no preference between the two diets (**Figure S1A**), suggesting that the added clozapine did not significantly change the taste of the control diet. Previous studies have shown that the metabolic effects of APDs are substantially more potent in female mice ^19,22,26^. Thus, we measured the chronic and acute effects of clozapine on energy metabolism in female C57BL/6 mice. Mass spectrometry analyses in clozapine-fed mice show that the concentration-to-dose ratio of plasma clozapine was comparable to that observed in human patients ^27^ (**Figure S1B**).

We found that mice fed the clozapine diet gained significantly more weight than those on the control diet over 12 weeks (**Figure 1A**). Nuclear magnetic resonance (NMR) analyses revealed that the excessive weight gain was primarily due to an increase in fat mass (**Figure 1B**), evident in both inguinal and gonadal white adipose tissues (iWAT and gWAT respectively) but not in the brown adipose tissue (BAT, **Figure 1C**). Consistent with this finding, hematoxylin and eosin (H&E) staining showed increased lipid accumulation in iWAT and liver of clozapine-fed mice (**Figures. S1C** and **S1D**). Additionally, clozapine-fed mice exhibited a deficit in glucose clearance in a glucose tolerance test (GTT, **Figure 1D**). These findings demonstrate that dietary supplementation of clozapine reproduces key symptoms of the drug-induced metabolic syndrome in humans.

**Figure 1.**
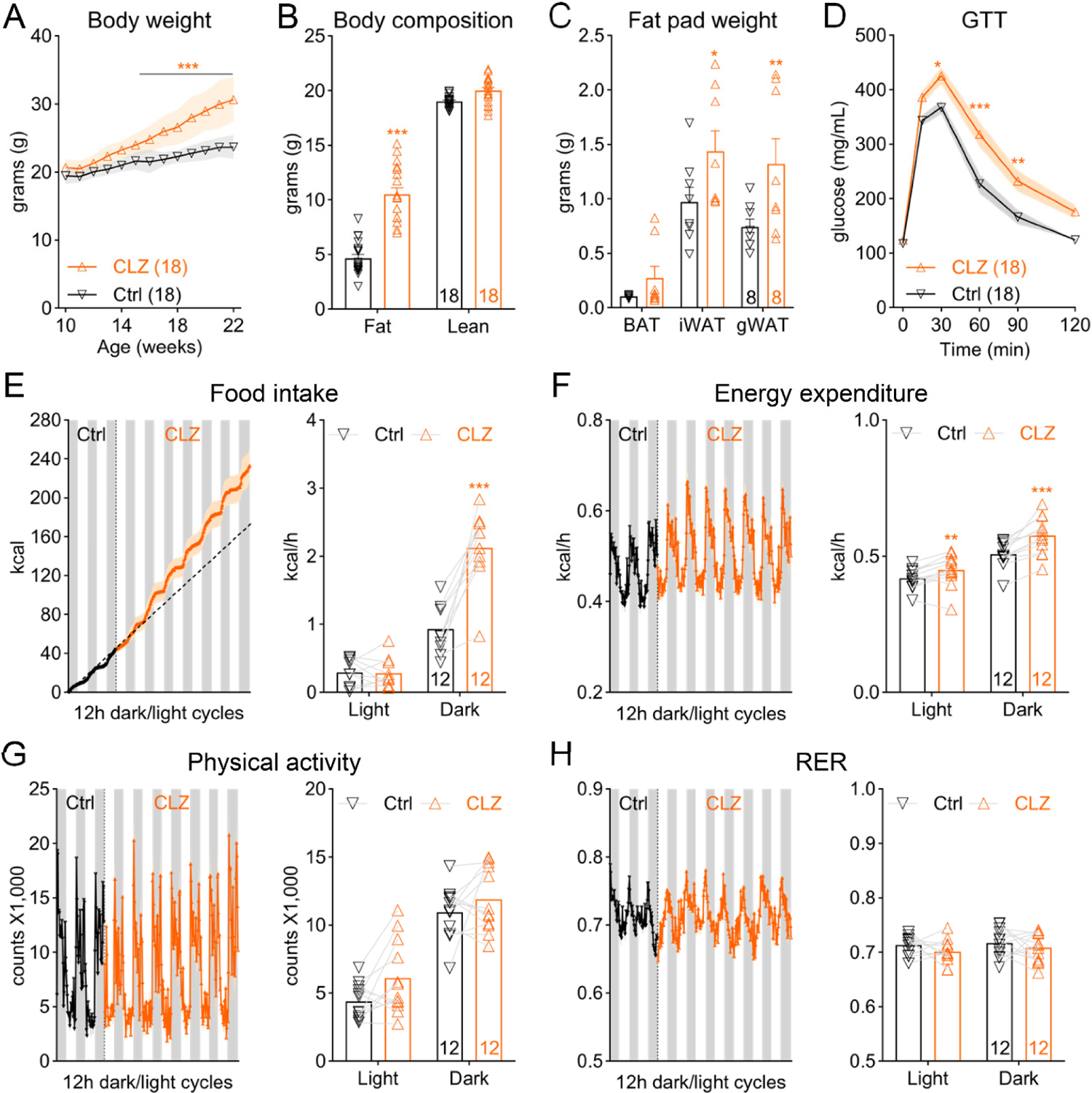
Clozapine exposure causes hyperphagia and obesity in female C57BL/6 mice. (A) Body weight of wild-type C57BL/6 mice fed either the control (Ctrl) or the clozapine (CLZ) diet; n=18, two-way ANOVA, F(1, 34)=49.53, p<0.001. (B) Body composition of mice after 12-week treatment of the Ctrl or CLZ diet; n=18, two-way ANOVA, F(1, 68)=39.7, p<0.001. (C) Fat pad weight in mice treated with the Ctrl or CLZ diet; n=8, two-way ANOVA, F(2, 42)=11.77, p=0.001. BAT, brown adipose tissue; iWAT, inguinal white adipose tissue; gWAT, gonadal white adipose tissue. (D) GTT after 12-week treatment of the Ctrl or CLZ diet; n=18, two-way ANOVA, F(1, 34)=10.57, p=0.003. (E-H) Left, traces of continuous measurement in metabolic cages; Right, summarized daily average (binned into 12-hour light and dark phases) before (black) and after (orange) the dietary switch; n=12, two-way ANOVA. (E) food intake, F(1, 9)=30.83, p<0.001; (F) energy expenditure, F(1, 11)=9.38, p=0.01; (G) physical activity, F(1, 11)=0.35, p=0.57; (H) RER, F(1, 11)=0.2, p=0.66. *p<0.05, **p<0.01, and ***p<0.001. Two-way ANOVA with Sidak’s *post hoc* tests. Data are presented as mean ± SEM.

### Clozapine-induced hyperphagia drives weight gain

To elucidate the mechanism behind clozapine-induced weight gain, we monitored individual mice’s energy intake and expenditure before and after clozapine treatment in a fully automated metabolic chamber system (TSE PhenoMaster). Mice were fed the control diet during acclimation and the first three days of the recording before transitioning to the clozapine diet for the next six days. Remarkably, mice quickly developed hyperphagia following the dietary switch, consuming more food during the dark phase of the day (**Figure 1E**). Additionally, energy expenditure increased in clozapine-fed mice (**Figure 1F**), while there was no difference in physical activity or respiratory exchange ratio (RER), an indicator of fuel utilization (**Figures 1G** and **1H**). Notably, despite the modest rise in energy expenditure (1.18 ± 0.23 kcal/day), it was insufficient to offset the significant increase in energy intake (11.75 ± 3.17 kcal/day), leading to a positive energy balance and weight gain in clozapine-fed mice.

### Clozapine treatment does not affect the canonical signaling of MC4R

Inhibition of MC4R^PVH^ neuron activity causes hyperphagia and obesity in mice ^20,21^. On murine hypothalamic slices, we demonstrated previously that risperidone acutely hyperpolarized MC4R^PVH^ neurons in an MC4R-dependent manner ^22^. To determine the impact of clozapine on these neurons, we injected adeno-associated viruses (AAV) expressing a Cre-dependent fluorescent calcium indicator (GCaMP8) into the PVH of *Mc4r-Cre* mice (**Figure 2A**). Fiber photometry experiments in freely moving mice reveal that an intraperitoneal (i.p.) dose of clozapine (3 mg/kg) reduced the activity of MC4R^PVH^ neurons *in vivo* (**Figure 2B**).

**Figure 2.**
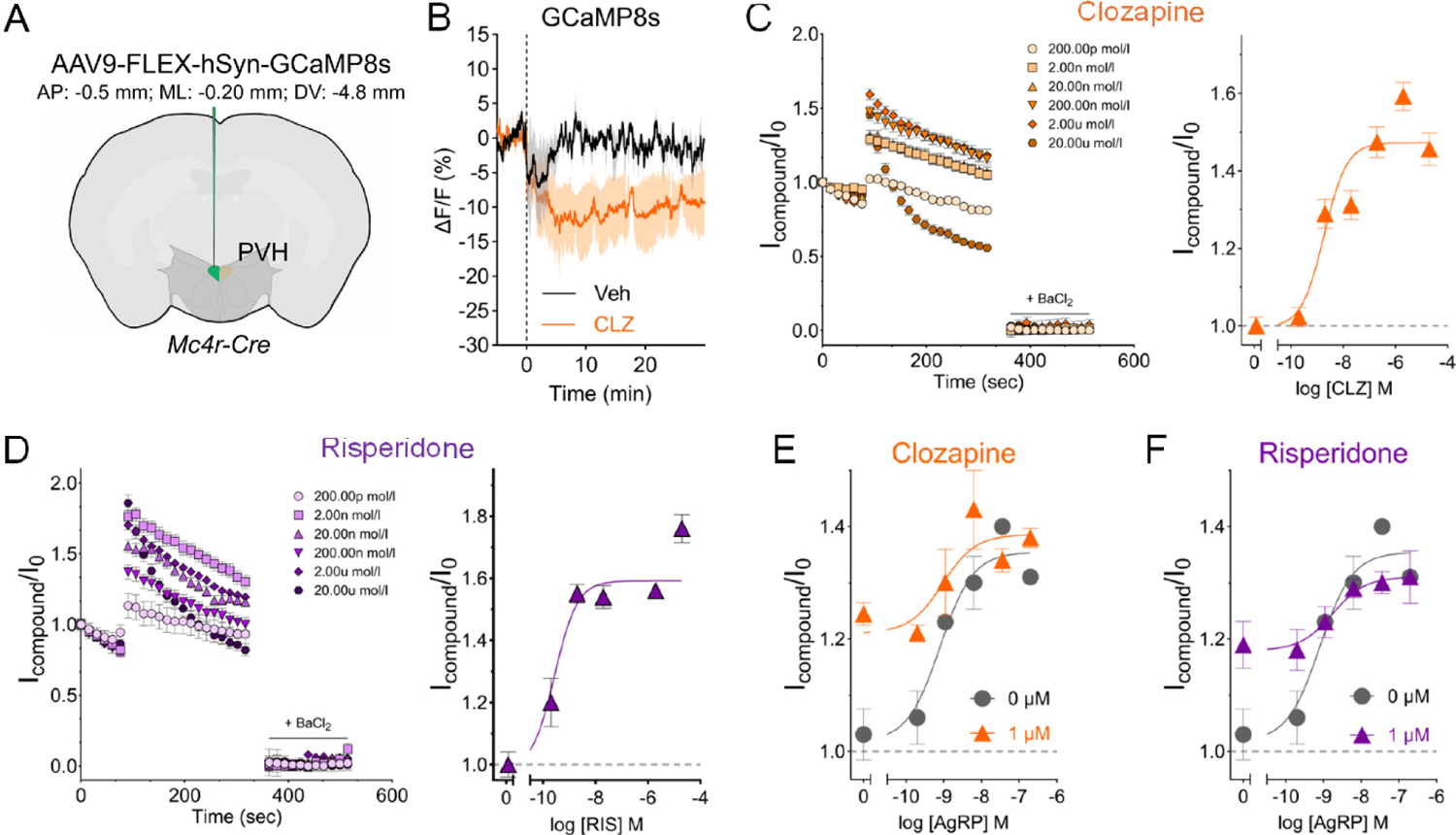
Clozapine and risperidone promote the Kir7.1 open state. (A) Schematic of the unilateral injections of AAV9-FLEX-GCaMP8s constructs into the PVH of *Mc4r-Cre* mice. (B) Calcium responses in MC4R^PVH^ neurons in living mice after an i.p. dose of vehicle (Veh) or clozapine (CLZ). (C and D) Left, whole-cell recordings of compound-evoked currents over baseline (I_Compound_/I_0_) time course; Right concentration-response curves for clozapine and risperidone in TREx HEK-293 cells expressing MC4R and Kir7.1. Data points represent the mean ± SEM of recordings obtained from 5 to 29 wells for each condition over three independent experiments. (E and F) AgRP concentration-response curves in the absence or presence of 1 μM clozapine (E) or risperidone (F). Data points represent the mean ± SEM. of recordings obtained from 7 to 13 wells for each condition over two independent experiments.

Upon binding with α-MSH or AgRP, MC4Rs depolarize or hyperpolarize MC4R^PVH^ neurons, respectively, via modulation of the activity of the potassium inward rectifier channel Kir7.1 ^18^. We tested the hypothesis that clozapine or other APDs may regulate MC4R^PVH^ neuron activity by competing for the orthosteric binding site of MC4R. In a radioligand binding assay, PF00446687 ^28^, a selective MC4R agonist, reduced the binding between MC4Rs and [^125^I], norleucine^4^, *D*-phenylalanine^7^ ⍺-MSH ([^125^I]-NDP-MSH, 100 pM, Ki = 199 ± 37 nM, mean ± S.E.M., n=3) in HEK 293 cells (**Figure S2A**). In contrast, risperidone and clozapine did not affect the radioligand binding, while clozapine only partially inhibited binding at the highest concentrations (10 and 50 µM) (**Figure S2A**). These findings suggest that the three APDs are unlikely to bind the orthosteric binding site under physiological conditions.

The canonical signaling cascade downstream of MC4R involves the activation of Gα_s_-coupled adenylyl cyclase, leading to an increase in intracellular adenosine monophosphate (cAMP). We investigated whether exposure to APD would alter the α-MSH-elicited cAMP responses in HEK 293 cells expressing MC4Rs and a genetic sensor for cAMP ^29^. We found that neither clozapine, risperidone, nor olanzapine (1 μM) affected α-MSH potency, efficacy, or the slope of the α-MSH-cAMP concentration-response curves, suggesting that these APDs do not influence the MC4R-Gα_s_ coupling or adenylyl-cyclase activation (**Figures S2B**, **S2C**, and **S2D**).

### Clozapine treatment increases MC4R-dependent Kir7.1 conductance

Activation of postsynaptic potassium currents contributes to risperidone-induced hyperpolarization of MC4R^PVH^ neurons ^22^. Notably, these neurons express the inwardly rectifying potassium channel Kir7.1 (encoded by the *kcnj13* gene) ^18,30^. Additionally, in HEK 293 cells co-expressing MC4R and Kir7.1, AgRP increases the Kir7.1 conductance in an MC4R-dependent manner ^17,18^. Using a high throughput automated patch clamp system (SyncroPatch 384PE, Nanion Technologies, Munich, Germany) ^31^, we investigated whether clozapine, risperidone, or olanzapine had a similar effect on MC4R-Kir7.1 coupling in HEK 293 cells. We measured the peak current ratios in the presence or absence of varying concentrations of APDs (I_Compound_/I_0_) to characterize the MC4R-Kir7.1-dependent potassium currents. Both clozapine and risperidone, at concentrations ranging from 0.2 nM to 20 μM, increased the inward potassium currents, with a pEC_50_ of 1.86 nM and 0.28 nM for clozapine and risperidone, respectively (**Figures 2C** and **2D**). In contrast, olanzapine did not increase potassium conductance but triggered channel closures (decreased potassium currents compared to the reference) at the highest concentration (10 μM, pIC_50_ of 1.95 μM, **Figure S3A**). Of note, treatment with α-MSH, AgRP ^17^, or individual APDs did not elicit potassium currents in HEK 293 cells expressing Kir7.1 alone (**Figure S4**). Additionally, we measured APDs’ effects on AgRP-induced Kir7.1 channel opening ^32^. We found that adding clozapine or risperidone (1 μM, respectively) increased the AgRP-induced basal MC4R-Kir7.1 coupling by 50%, with no changes in the AgRP potency, efficacy, or concentration-response curve slopes (**Figures 2E** and **2F**). In comparison, olanzapine (1 μM) did not affect the AgRP-induced opening of Kir7.1 (**Figure S3B**).

### *Kir7*.1 in *Mc4r-Cre* neurons is necessary for clozapine-induced weight gain

To determine the role of *Kir7.1* in clozapine-induced weight gain, we generated *Kir7.1^fl/fl^; Mc4r-Cre +/-* mice (designated hereafter as *Kir7.1^Mc4r KO^* mice) in which *Kir7.1* was selectively deleted in *Mc4r-Cre*-expressing neurons. Like wild-type C57BL/6 mice, *Kir7.1^fl/fl^* mice fed the clozapine diet gained significantly more weight and adiposity (**Figures S5A** and **S5B**). In contrast, clozapine’s effect on body weight and fat was diminished in *Kir7.1^Mc4r KO^* mice (**Figures 3A** and **3B**). Despite the lack of weight gain, clozapine-induced glucose intolerance persisted in *Kir7.1^Mc4r KO^* mice (**Figure 3C**). On the other hand, feeding mice an olanzapine diet ^33^ led to significant weight gain and fat accumulation in both *Kir7.1^Mc4r KO^* (**Figures 3D** and **3E**) and *Kir7.1^fl/fl^* (**Figures S5C** and **S5D**) mice, suggesting that *Kir7*.1 in *Mc4r-Cre* neurons is not essential for the obesogenic effect of olanzapine.

**Figure 3.**
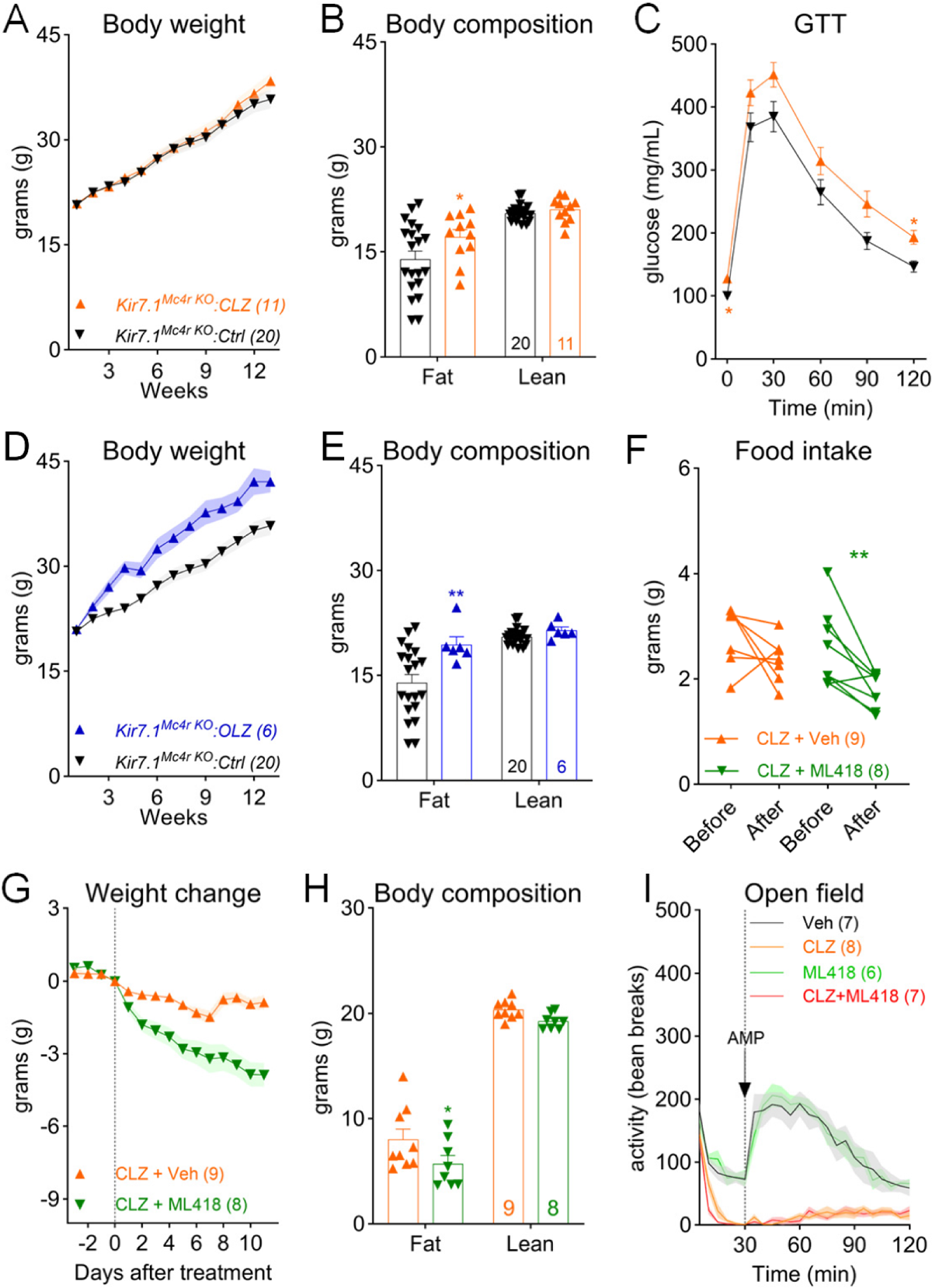
*Kir7.1* in *Mc4r-Cre* neurons is necessary for clozapine-induced weight gain (A) Body weight of *Kir7.1^Mc4r KO^* mice fed either the control (Ctrl) or the clozapine (CLZ) diet; n=11-20, two-way ANOVA, F(1, 29)=0.29, p=0.59. (B) Body composition of mice after 12-week treatment of the Ctrl or CLZ diet; n=11-20, two-way ANOVA, F(1, 29)=3.24, p=0.08. (C) GTT after 12-week treatment of the Ctrl or CLZ diet; n=11-20, two-way ANOVA, F(1, 32)=6.08, p=0.02. (D) Body weight of *Kir7.1^Mc4r KO^* mice fed either the control (Ctrl) or the olanzapine (OLZ) diet; n=6-20, two-way ANOVA, F(1, 24)=9.38, p=0.005. (E) Body composition of mice after 12-week treatment of the Ctrl or CLZ diet; n=11-20, two-way ANOVA, F(1, 24)=5.77, p=0.02. (F) Changes in daily food intake in clozapine-fed C57BL/6 mice treated with ML418 or vehicle (Veh); paired t-test, **p<0.01. (G) Changes in body weight before and after the treatment of ML418 or Veh; n=8-9, two-way ANOVA, F(1, 15)=18.37, p<0.001. (H) Body composition in clozapine-fed mice after treatment of ML418 or Veh; n=8-9, two-way ANOVA, F(1, 15)=4.78, p=0.05. (I) Open field test measuring amphetamine-induced hyperlocomotion in mice pretreated with Veh, CLZ, ML418 or CLZ + ML418; n=6-8, two-way ANOVA, F(69, 552)=10.31, p<0.0001. *p<0.05 and **p<0.01. Two-way ANOVA with Sidak’s *post hoc* tests. Data are presented as mean ± SEM.

### Blockade of Kir7.1 reduces clozapine-induced hyperphagia

Next, we investigated whether pharmacological blockade of the channel Kir7.1 could mitigate clozapine-induced overeating and weight gain using ML418, a small molecular channel blocker with significant specificity for Kir7.1 ^34^. To this end, we fed C57BL/6 mice the clozapine diet for 12 weeks to induce hyperphagia and obesity. We then treated them with a daily dose of either ML418 (5 mg/kg, i.p.) or vehicle while maintaining the mice on the clozapine diet. We found that ML418 treatment (5 mg/kg/d) reduced hyperphagia in clozapine-fed mice (**Figure 3F**), resulting in a significant decrease in body weight after the 10-day treatment period (**Figure 3G**). Consistent with this finding, NMR analyses showed a selective reduction in the fat mass of these mice (**Figure 3H**).

Additionally, we investigated whether co-treatment with ML418 affected clozapine’s psychotropic effect using a rodent model of schizophrenia ^35^. It is well established that amphetamine induces schizophrenia-like psychotic episodes in healthy human subjects ^36^ and exacerbates symptoms in schizophrenic patients ^37^. Similarly, administration of amphetamine (2.5 mg/kg, i.p.) induced hyperactivity in C57BL/6 mice, a response significantly attenuated by pretreatment with clozapine (**Figure 3I**). Importantly, we found that co-treatment with ML418 did not diminish clozapine’s ability to suppress amphetamine-induced hyperlocomotion in an open-field test (**Figure 3I**).

## DISCUSSION

### A mouse model of clozapine-induced hyperphagia

APD-induced weight gain and metabolic syndrome affect millions of patients. However, the difficulty of modeling their metabolic effects in lab mice has impeded relevant mechanistic studies. In this study, we developed a clozapine diet that causes excessive weight gain in female C57BL/6 mice. Despite a lower dose, dietary supplementation of clozapine resulted in more pronounced metabolic disturbances than those induced via other routes ^24,25^. The mechanism behind the increased efficacy remains less clear. Clozapine may be more stable when mixed in the diet ^38^. Moreover, oral delivery permits drug interactions with the microbiota, which have been implicated in APD-induced weight gain ^39^.

The new mouse model enables us to determine the pathophysiology underlying clozapine-induced weight gain. Metabolic chamber analyses revealed that clozapine treatment profoundly alters energy intake and expenditure in mice. We found that hyperphagia develops shortly after clozapine exposure. Others have shown that clozapine treatment promotes thermogenesis by enhancing the activities of the sympathetic nerves that innervate the brown adipose tissue ^40^. Consistent with this finding, energy expenditure increases after clozapine treatment, which protects against weight gain. However, the modest rise in energy expenditure is insufficient to counterbalance the extra energy intake, suggesting that drug-induced hyperphagia is the driving force behind weight gain.

### APDs target the central melanocortin system to cause weight gain

Various neural pathways have been implicated in APDs’ metabolic effects ^9 10,11 12 26^. Given their unique binding profiles, individual APDs likely engage distinct brain circuits to alter energy metabolism. However, emerging evidence suggests that some commonly used APDs may converge on central melanocortin neurons to induce hyperphagia and weight gain ^19,22^.

In mammals, MC4R signaling in the brain is a crucial regulator of energy homeostasis. In particular, MC4R in PVH neurons plays a pivotal role in regulating food intake ^22,41^ while modulating energy expenditure in other brain regions ^42^. Human studies have linked common variants near MC4R with APD-induced weight gain ^43^. Moreover, we have demonstrated that the obesogenic effects of olanzapine and risperidone are diminished in mice lacking MC4R in PVH neurons ^22^.

APDs may disturb MC4R function via different means. For example, olanzapine exposure increases the expression of the orexigenic neuropeptide AgRP, which acts via the MC4Rs ^19^. Meanwhile, APDs can promote feeding by directly reducing the activity of MC4R^PVH^ neurons. Silencing these neurons in adult mice is sufficient to cause hyperphagia and obesity ^20,21^. Indeed, we have demonstrated that risperidone acutely inhibits MC4R^PVH^ neurons in hypothalamic brain slices ^22^. The current study shows that clozapine has a similar effect on these neurons *in vivo*. These findings suggest that MC4R/MC4R^PVH^ neurons may be a common target for different APDs to cause weight gain.

### MC4R-Kir7.1 coupling in APD-induced hyperphagia

The mechanism whereby APDs inhibit MC4R^PVH^ neurons was largely unknown prior to our study. Alpha-MSH and AgRP can directly depolarize or hyperpolarize MC4R^PVH^ neurons, respectively ^18^. This raises the possibility that APDs might interfere with binding between MC4R and its agonists. However, our findings reveal that none of the three APDs tested compete for the orthosteric binding site of MC4R in a radioligand binding assay. Similarly, they do not affect the Gαs-coupled adenylyl-cyclase activation or intracellular cAMP responses.

AgRP-induced hyperpolarization of MC4R^PVH^ neurons is partly attributed to an increase in postsynaptic Kir7.1 conductance. Moreover, this effect is independent of the canonical Gαs signaling but relies on the functional coupling between MC4R and Kir7.1 ^18^. The observation of risperidone-evoked potassium currents in MC4R^PVH^ neurons ^22^ suggests that APDs may inhibit these neurons by activating Kir7.1. Indeed, whole-cell patch clamp recordings in HEK 293 cells demonstrate that risperidone and clozapine dose-dependently promote the opening state of the Kir7.1. Notably, this effect depends on MC4Rs, as neither drug induces potassium currents in cells expressing Kir7.1 alone. Moreover, both drugs enhance AgRP-induced Kir7.1 conductance without affecting the potency or affinity of AgRP. We suspect that clozapine and risperidone might allosterically modulate the receptor-channel complex formation, coupling, or signaling modality in a G-protein-independent manner. Moreover, these findings suggest that clozapine induces hyperphagia and weight gain by promoting MC4R-dependent Kir7.1 conductance in MC4R^PVH^ neurons. Supporting this hypothesis, clozapine treatment fails to increase body weight in mice lacking Kir7.1 specifically in *Mc4r-Cre* neurons.

Surprisingly, olanzapine does not affect MC4R-Kir7.1 coupling, and deleting Kir7.1 in *Mc4r-Cre* neurons does not prevent olanzapine-induced weight gain. These findings highlight the divergent mechanisms through which APDs induce weight gain. Olanzapine might still inhibit MC4R^PVH^ neurons via MC4R-Kir7.1 independent mechanisms. Alternatively, it may indirectly perturb MC4R function by increasing AgRP expression. Indeed, olanzapine’s effect on food intake and body weight is abolished in *Agrp*-deficient mice ^19^.

### Targeting MC4R-Kir7.1 coupling to mitigate APD-induced weight gain

Therapies aimed at addressing APD-induced metabolic syndromes have been inadequate. Our findings suggest that MC4R-Kir7.1 coupling represents a potential new therapeutic target for managing clozapine and risperidone-induced weight gain. Furthermore, when developing new antipsychotic medications, considerations should be made regarding their potential effects on MC4R-Kir7.1 coupling to minimize side effects.

In contrast to AgRP, α-MSH reduces MC4R-dependent Kir7.1 conductance ^18^. As anticipated, setmelanotide, an MC4R agonist and weight-loss medication ^44^, counteracted risperidone-induced hyperpolarization of MC4R^PVH^ neurons and reversed obesity in risperidone-fed mice. Likewise, we demonstrate that the Kir7.1 blocker, ML418, alleviates hyperphagia and weight gain in mice treated with clozapine. Importantly, co-administration of neither setmelanotide nor ML418 diminished risperidone or clozapine’s ability to suppress schizophrenia-like hyperlocomotion in mice. These findings suggest that the psychotropic and metabolic effects of clozapine and risperidone may involve distinct neural pathways and thus can be targeted separately.

### Limitations of the study

Our study reveals the molecular mechanisms underlying olanzapine-induced hyperphagia and suggests MC4R-Kir7.1 signaling as a novel target for mitigating clozapine and risperidone-induced weight gain. It remains to be determined whether clozapine or risperidone directly binds to the MC4R-Kir7.1 complex to modulate its activity. Moreover, potential drug effects on MC4Rs in other hypothalamic and extra-hypothalamic neurons were not studied.

Our findings are relevant to human diseases; however, they are derived from studies conducted in laboratory mice. Therefore, further investigation is necessary to ascertain whether these findings can be translated into potential pharmacotherapies targeting the MC4R-Kir7.1 pathway to mitigate weight gain in human patients.

## STAR METHODS

- KEY RESOURCES TABLE
- RESOURCE AVAILABILITY

- Lead contact
- Further information and requests for resources and reagents should be directed to and will be fulfilled by the lead contact, Chen Liu
- Materials availability
- We have open-sourced the ingredients for the clozapine diet described in this paper (please refer to Table S1). The clozapine (#D21070104) and control (D09092903) diets are available at Research Diets Inc.
- Data and code availability
- The lead contact will share data reported in this paper upon request. This paper does not report original code. Any additional information required to reanalyze the data reported in this paper is available from the lead contact upon request.

## EXPERIMENTAL MODEL AND SUBJECT DETAILS

All mice were housed in a temperature-controlled room in the UT Southwestern Medical Center animal facility with a 12-hour light/12-hour dark cycle (lights on at 06:00 am, lights off at 6:00 pm). Unless otherwise noted, mice were provided standard chow (No. 2016; Harlan Teklad) and water ad libitum. C57BL/6 mice were purchased from the Jackson Laboratory (#000664). The *Kir7.1^fl/f^* mouse line ^45^ was generously provided by Dr. Graeme Mardon at Baylor College of Medicine. *Mc4r-Cre* +/-; *Kir7.1^fl/fl^* and *Mc4r^fl/fl^* littermates were maintained on a C57BL/6 background. The Institutional Animal Care and Use Committee of the University of Texas Southwestern Medical Center approved all mouse experiments.

## METHOD DETAILS

### Diets

All diets were prepared by ResearchDiets Inc. and shared the same macronutrients, ingredient composition, and energy density (4.76 kcal/gram). Clozapine (MedChemExpress, Cat# HY-14539) was mixed into a control diet D09092903 to make the clozapine diet D21070104 (50 mg/kg). Similarly, olanzapine (ChemPacific, Cat# 356941G) was mixed into the same control diet, D09092903, to make the olanzapine diet as we described previously ^33^.

### Metabolic phenotype analysis

We used a magnetic-resonance whole-body composition analyzer (EchoMRI) to analyze the mouse’s body composition (fat mass, lean mass, and water content). For chronic studies, body weight and food intake were measured weekly. The acute effects of clozapine on energy intake and expenditure were measured using an indirect calorimetric system (TSE PhenoMaster, Germany) in the Metabolic Phenotyping Core of UT Southwestern Medical Center.

### Radioligand binding experiments

Radioligand binding experiments used a HEK-293 (ATCC CRL-1573™) stable cell line expressing the human MC4R. The cells were grown in 150 mm culture dishes to prepare crude membranes until they reached 90-95% confluency. TrypLE Express (1x) cell dissociation reagent (Thermo Fisher Scientific) was used to detach the cells from the growth vessels. Afterward, trypsin was neutralized with an equal volume of complete growth media. The cells were pelleted at 200 × g for 5 min, washed twice with phosphate-buffered saline (PBS, and resuspended in “Membrane Extract Buffer” (ME-buffer). This buffer contained 50 mM Tris-HCl (pH 7.4), 10 mM MgSO_4_, 1.5 mM CaCl_2_, 0.5 mM EDTA, and cOmplete™ EDTA-free protease inhibitor cocktail (Roche, Basel Switzerland). The initial cell pellet was homogenized using a 15 mL Dounce homogenizer. Homogenates were centrifuged at 35,000 × g for 20 min, and the process was repeated twice. The final pellet was resuspended with 10× v/w of ME-buffer supplemented with 10% sucrose, and aliquots were stored at −80℃. The total protein content from the crude membrane extracts was determined using the bicinchoninic acid assay ^46^ with a commercially available kit (Pierce™ BCA Protein Assay Kit, Thermo Fisher Scientific).

Equilibrium competition binding experiments were performed with 4 µg of total protein from the crude membrane extract. [^125^I]-[Nle^4^, DPhe^7^]-ɑ-MSH (NDP-MSH, specific activity: 2200 Ci/mmol)) was used as the radioligand tracer at a concentration of 100 pM. This radioligand concentration resembles the equilibrium dissociation constant (K_D_) determined by ligand saturation experiments. Varying concentrations of the indicated antipsychotic drugs (APD) or PF-00446687, an MC4-R-specific agonist, were incubated with the crude membrane extracts in a final volume of 200 µl of Binding Assay buffer (BA-buffer: 25 mM HEPES pH 7.4, 1 mM MgSO_4_, 1 mM CaCl_2_, 0.2% bovine serum albumin (BSA), and cOmplete™ EDTA-free protease inhibitor cocktail). APD and crude membranes were preincubated for 16 hours at room temperature before adding radioligand to ensure equilibrium. The assay was incubated for an additional 120 min in the presence of the radioligand. Bound [^125^I]-NDP-MSH was rapidly separated from the free ligand by filtration through GF/B glass fiber filters (Brandel, Gaithersburg, MD) using a cell harvester (Brandel) manifold. During filtration, the filters were washed five times rapidly with cold 25 mM Tris-HCl pH 7.4. IC_50_ values were derived from nonlinear regression fitting the data to a sigmoid function model. Ki values were derived by applying the Cheng and Prusoff transformation ^47^.

### Cellular cAMP responses

A HEK-293 (ATCC CRL-1573™) stable cell line expressing the human MC4-R and a split luciferase cAMP genetic sensor based on the circularized fusion of firefly luciferase and protein kinase A regulatory cAMP binding domain (clone 22F, Promega, Madison WI) was used to determine the α-MSH-elicited cAMP responses in live cells as previously described ^32^. The cell line identity was verified by qPCR and MC4-R subtype-specific oligonucleotides ^48^.

Briefly, twenty-four hours before the cAMP response recordings, the cells were plated in poly-D-lysine coated clear-bottom black wall 384-well plates (Corning Inc. Corning, NJ) at a density of 20,000 cells per well in growth medium (Dulbecco’s Modified Eagle Medium (DMEM) supplemented with 4.5 g/L D-glucose, 110 mg/L sodium pyruvate, 10% fetal bovine serum, 1× Antibiotic-Antimycotic™ cocktail (Thermo Fisher Scientific, Waltham, MA), 200 μg/mL hygromycin B, and 700 μg/mL Geneticin™. Two hours before the start of the experiment, the growth medium was replaced with CO_2_ Independent Medium™ (Thermo Fisher Scientific), containing 4% D-luciferin (Promega). The luciferase substrate permeated the cells for 120 min at 37 ℃. Intracellular cAMP levels were measured using an FDSS 7000EX Functional Drug Screening System (Hamamatsu Photonics, Hamamatsu, Japan). To test the effect of APD on the α-MSH concentration-response profile, the in-line addition of 1 μM (final concentration) clozapine, risperidone, or olanzapine was performed and allowed to reach equilibrium for 12 minutes. This was followed by adding 10 µL of varying 3× concentrations of α-MSH or vehicle. Luminescence responses were measured for an additional 12 min. The agonist responses were normalized to the α-MSH maximum response without APD. The data were fit to a four-parameter sigmoid model, constraining the slope to a range of 0 to 2 using GraphPad Prism.

### Histology

Mouse adipose tissue was fixed in 4% paraformaldehyde overnight and then dehydrated through a series of ethanol baths with ascending concentrations (up to 100%). The Histo Pathology Core of the University of Texas Southwestern Medical Center performed paraffin embedding, sectioning, and H&E staining.

### Automated Patch Clamp

Patch clamp recordings were conducted using a SyncroPatch 384PE system from Nanion Technologies (Munich, Germany) at the Center for Chemical Genomics at the University of Michigan. Recordings were carried out in the whole cell voltage clamp configuration employing medium resistance four-hole NPC-384 chips. Data acquisition and analysis were performed with PatchControl 384 (for data acquisition) and DataControl 384 (for data analysis), both from Nanion Technologies. The experimental setup utilized an inducible T-REx HEK293 cell line (Thermo Fisher Scientific) expressing MC4-R and the inwardly rectifying K^+^ channel Kir7.1 mutant M125R. Cells were centrifuged at 200*g* for 5 minutes, resuspended in an external solution at a density of 500,000 cells/mL, and added to the instrument cell hotel. The cell hotel was maintained at 10 °C with 200 rpm agitation until the start of the experiment. The external recording solution comprised (in mM): NaCl 140, KCl 4, CaCl_2_ 2, MgCl_2_ 1, HEPES 10, glucose 5; pH 7.4 (NaOH, ∼300 mOsm). Cell sealing was accomplished in a modified external solution containing 6 mM CaCl_2_. The internal electrode solution included (in mM): KF 110, KCl 10, NaCl 10, HEPES 10, EGTA 10; pH 7.2 (KOH, ∼280 mOsm).

To assess MC4-R-dependent Kir7.1 channel closures or activations induced by antipsychotic drugs (APD), a two-stage voltage protocol was applied. This protocol consisted of a hyperpolarization step to −140 mV for 200 ms, followed by a ramp (−1.2 V/s) from +20 to −180 mV for 500 ms (holding potential −60 mV), repeated every 5 seconds. Peak Kir7.1 inward current amplitude during the hyperpolarizing step was measured and normalized to the remnant currents after adding 10 mM BaCl_2_ (full block). Antipsychotic drugs (APD) were delivered at various concentrations across the NPC-384 chip, followed by a Ba^2+^ full channel block added to each well to determine non-K^+^ channel currents, as the M125R Kir7.1 regains sensitivity for Ba^2+^ compared to the relatively insensitive wild-type Kir7.1. Responses were further normalized by determining the ratios of peak currents in the presence or absence of APDs (I_compound_/I_0_). Concentration-response profiles were determined, and EC_50_ or IC_50_ values for activating or inhibiting APD responses were calculated. A seal quality assessment for each well was performed using the DataControl 384 application, excluding rundown wells, with a positive patch rate of 87%.

### GTT

For the glucose tolerance test (GTT), mice were fasted for 7 hours with water provided *ad libitum* from 8 am on the experimental day. During GTT, blood glucose levels were monitored at 0, 15, 30, 60, 90, and 120 minutes after an i.p. dose of glucose (Dextrose) (1.0 g/kg body weight). Blood glucose was taken from the tail vein and analyzed using a Glucometer (Johnson & Johnson).

### Amphetamine-induced hyperlocomotion

The Rodent Behavior Core performed the amphetamine-induced hyperlocomotion in the Peter O’Donnell Jr. Brain Institute at UT Southwestern Medical Center. Briefly, C57BL/6 mice were habituated for the open field test with daily single i.p. saline injections for three days. On the third day of habituation, all mice were given D-amphetamine (2.5 mg/kg) 30 min after the saline injection. Following the habituation, all mice were randomly divided into four groups: 1) mice in the saline group received an i.p. dose of saline; 2) Mice in the ML418 group received an i.p. dose of ML418 (5 mg/kg); 3) mice in the CLZ group received an i.p. dose of clozapine (3 mg/kg) 4) mice in the ML418 + CLZ group received both clozapine (3 mg/kg) and ML418 (5 mg/kg). Locomotor activity (bean breaks) was immediately recorded after the first injection. An i.p. dose of D-amphetamine (2.5 mg/kg) was administered 30 min after the first injection.

### Quantification and statistical analyses

Sample sizes were chosen to reliably measure experimental parameters while remaining in compliance with ethical guidelines for minimizing animal use, and they were similar to those reported in previous publications. All the virus expression and optic fiber implant placement were verified by *post hoc* histology. Replicate information is indicated in the figure legends. All results are presented as mean ± SEM and analyzed using statistical tools implemented in Prism (GraphPad, version 10). Statistical analyses were performed using the Student’s *t*-test and regular one-way or two-way analysis of variance (ANOVA). Differences with *P* < 0.05 were considered to be significant. *P* < 0.05 (*), *P* < 0.01 (**), and *P* < 0.001 (***).

## SUPPLEMENTAL INFORMATION

Supplemental information can be found online at

## ACKNOWLEDGMENTS

CL was supported by the US NIH grants R01 DK114036, DK130892, and DK136592. LL was supported by a postdoctoral fellowship (23POST1019715) and a Career Development Award (24CDA1257999) from the American Heart Association. RDC was supported by R01 DK070332. The Nanion Synchropatch used for patch clamp recordings was supported by NIH S10OD025203. We thank members of the UTSW Metabolic Phenotyping Core, which was supported by the UTSWNORC grant under NIDDK/NIH award number P30DK127984. We thank Drs. Noelle Williams and Xiaoyu Wang from UTSW Preclinical Pharmacology Core. The behavior testing presented in this publication was performed by the staff of the Rodent Behavior Core, which is supported by the UT Southwestern Peter O’Donnell Jr. Brain Institute.

## AUTHOR CONTRIBUTIONS

Author contributions: LL, CCH, LEG, RDC, and CL designed the experiments. LL, CCH, LEG, BX, S, and SGB collected data. LL, CCH, LEG, RDC, and CL analyzed the data and wrote the manuscript.

---

**Figure S1.**
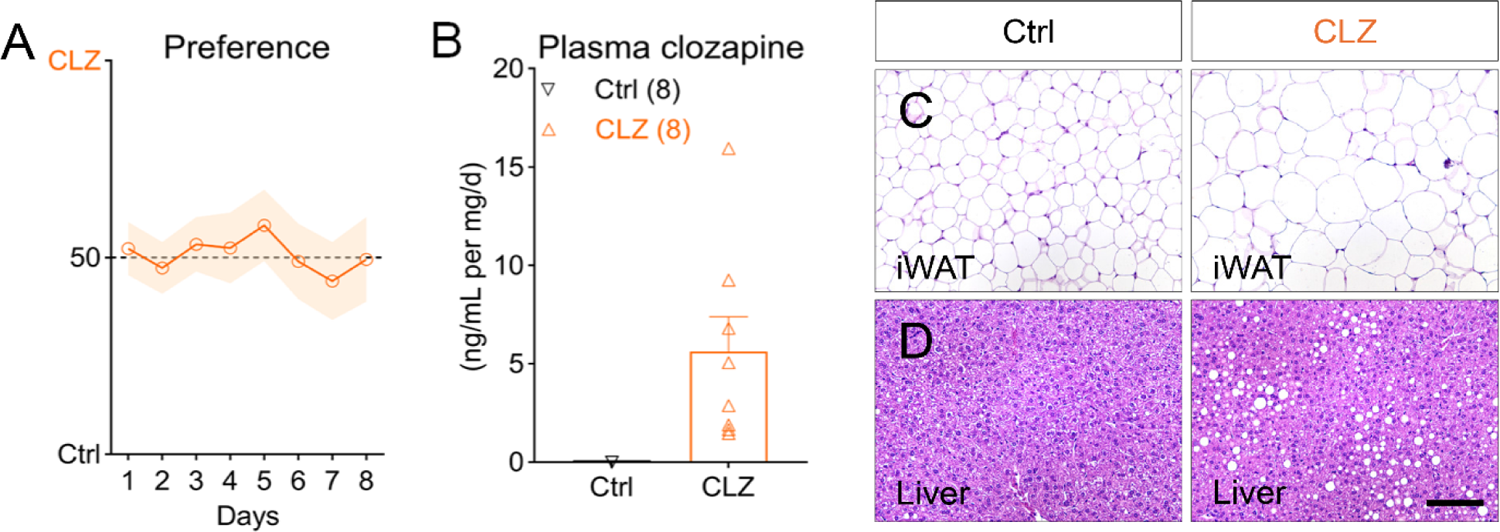
Chronic clozapine treatment increases lipid accumulation. (A) Ratio of daily intake of the CLZ diet (50 mg/kg) over that of the Ctrl diet when both diets were available in the cage for 8 days, n=10. (B) Clozapine concentration-to-dose ratio in plasma based on an estimated daily intake of 4 g of clozapine diet, n=8. (C and D) H&E staining in mice after 12-week treatment of the Ctrl or CLZ diet. Scale bar is 100 µm. Data are presented as mean ± SEM.

**Figure S2.**
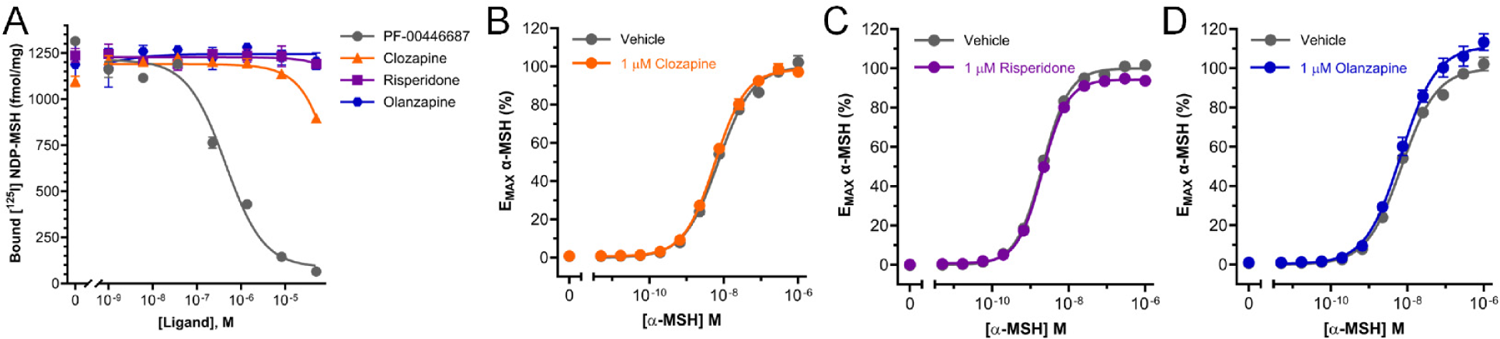
Clozapine, risperidone, and olanzapine do not interact with the MC4R orthosteric binding site. (A) [^125^I]-NDP-MSH (100 pM) ligand binding (fmol/mg of total crude homogenate proteins) in the presence of varying concentrations (1 nM to 1 μM) of PF-00446687, clozapine, risperidone, or olanzapine. The data represent the mean ± SEM from triplicate data points fit to a three-parameter sigmoid model. One out of two independent experiments is shown. (B-D) α-MSH cAMP concentration-response curves in the presence of 1 μM clozapine (B), risperidone (C), or olanzapine (D). The data represent the mean ± SEM. from triplicate data points fit to a three-parameter sigmoid model. One out of two independent experiments is shown.

**Figure S3.**
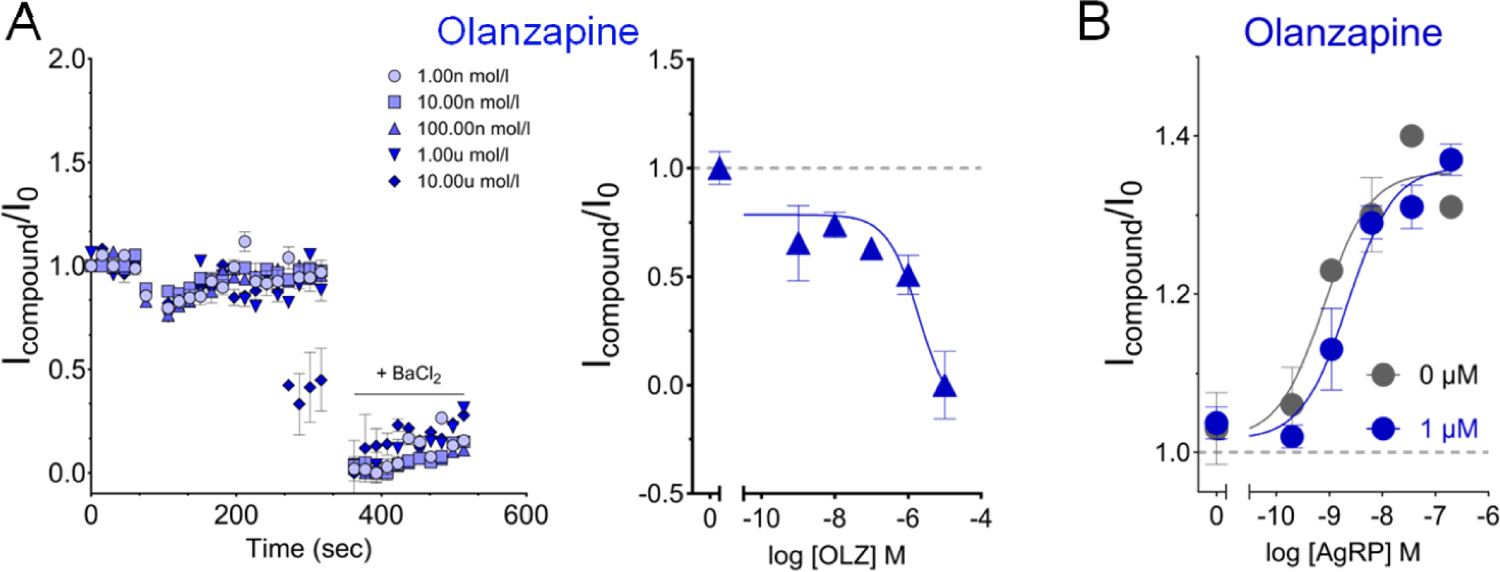
Olanzapine promotes the channel closed state. (A) Left, whole-cell recordings of compound-evoked currents over baseline (I_Compound_/I_0_) time course, Right, concentration-response curves for olanzapine in TREx HEK-293 cells expressing MC4R and Kir7.1. Data points represent the mean ± SEM of recordings obtained from 7 to 10 wells for each condition over two independent experiments. (B) AgRP concentration-response curves in the absence or presence of 1 μM olanzapine. Data points represent the mean ± SEM. of recordings obtained from 9 to 11 wells for each condition over two independent experiments.

**Figure S4.**
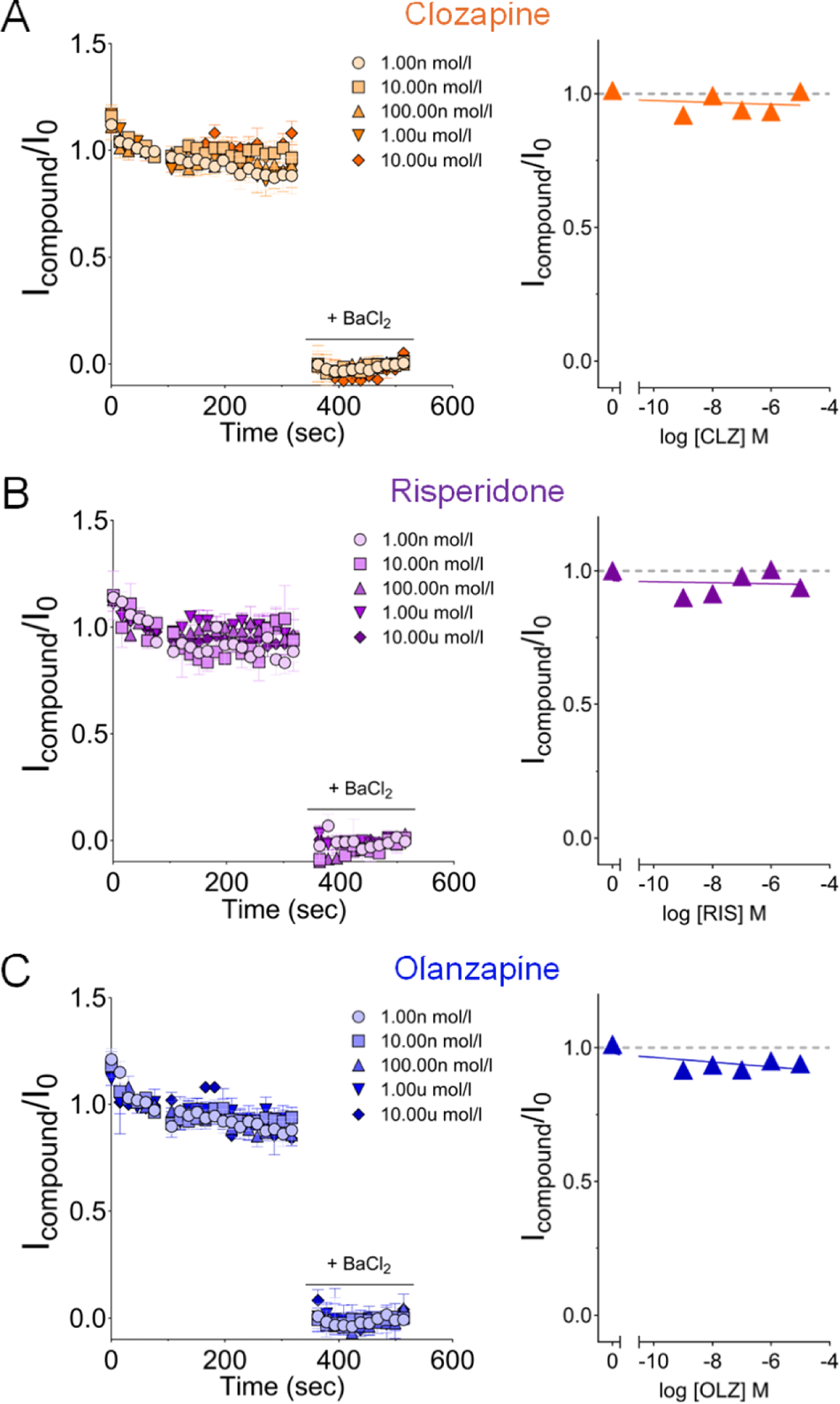
APDs have no direct effect on the Kir7.1 function. (A-C) Whole-cell recordings of compound-evoked currents over baseline (I_Compound_/I_0_) time course (left panels) and concentration-response curves (right panels) in the presence of the indicated concentrations of clozapine (A), risperidone (B), or olanzapine (C). The data points represent the mean ± SEM of recordings obtained from 15 wells for each condition over two independent experiments.

**Figure S5.**
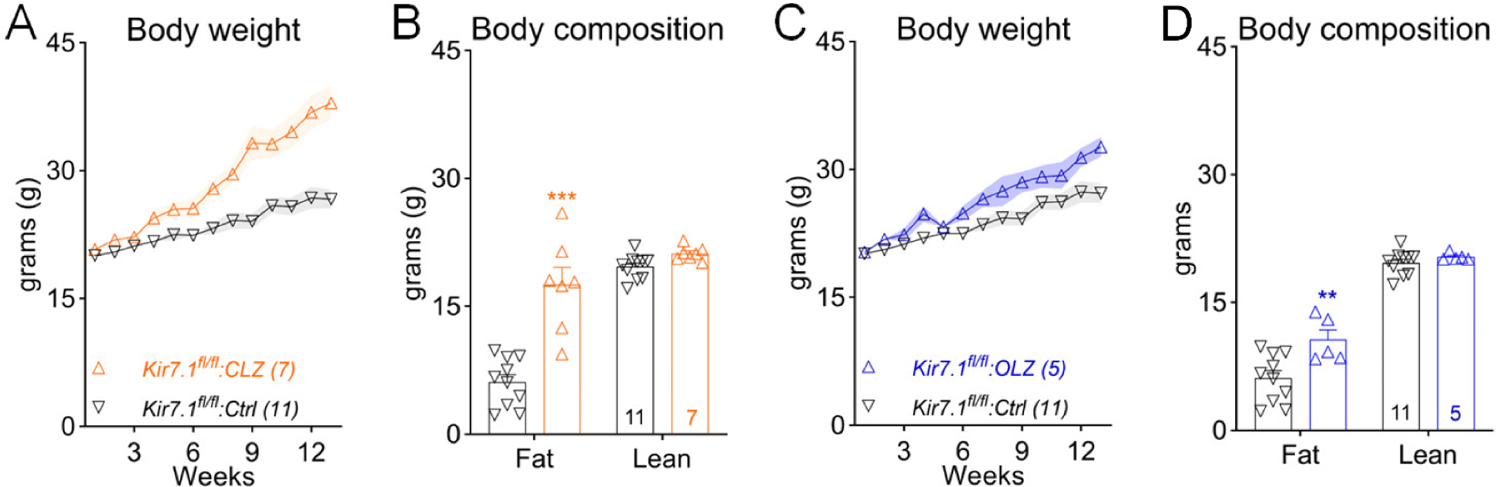
Chronic treatment with clozapine or olanzapine causes obesity in *Kir7.1^fl/fl^* mice (A) Body weight of *Kir7.1^fl/fl^* mice fed either the control (Ctrl) or the clozapine (CLZ) diet; n=7-11, two-way ANOVA, F(1, 15)=17.7, p<0.001. (B) Body composition of mice after 12-week treatment of the Ctrl or CLZ diet; n=7-11, two-way ANOVA, F(1, 15)=31.1, p<0.001. (C) Body weight of *Kir7.1^fl/fl^* mice fed either the control (Ctrl) or the olanzapine (OLZ) diet; n=5-11, two-way ANOVA, F(1, 14)=4.46, p=0.05. (D) Body composition of mice after 12-week treatment of the Ctrl or OLZ diet; n=5-11, two-way ANOVA, F(1, 13)=7.55, p=0.02. **p<0.01 and ***p<0.001 Two-way ANOVA with Sidak’s *post hoc* tests. Data are presented as mean ± SEM.

**Table S1.** Macronutrient composition and ingredients of the control and clozapine diets.

| **Diets** | **Control** |  | **Clozapine** |  |
| --- | --- | --- | --- | --- |
|  | **45 kcal% fat** |  | **45 kcal% fat** |  |
| **Composition** | gm% | kcal% | gm% | kcal% |
| Protein | 25 | 21 | 25 | 21 |
| Carbohydrate | 41 | 34 | 41 | 34 |
| Fat | 24 | 45 | 24 | 45 |
| Total |  | 100.0 |  | 100.0 |
| kcal/gm | 4.77 |  | 4.76 |  |
| **Ingredient** | **gm** | **kcal** | **gm** | **kcal** |
| Casein | 200 | 800 | 200 | 800 |
| DL-Methionine | 3 | 12 | 3 | 12 |
| Corn Starch | 50 | 200 | 50 | 200 |
| Maltodextrin | 100 | 400 | 100 | 400 |
| Sucrose | 173.8 | 695 | 173.8 | 695 |
| Cellulose | 50 | 0 | 50 | 0 |
| Corn Oil | 50 | 450 | 50 | 450 |
| Lard | 145 | 1305 | 145 | 1305 |
| Mineral Mix S10001 | 35 | 0 | 35 | 0 |
| Vitamin Mix V10001 | 10 | 40 | 10 | 40 |
| Choline Bitartrate | 2 | 0 | 2 | 0 |
| **Clozapine** | 0 | 0 | **0.041** | **0** |
| **Total** | **818.9** | **3902** | **818.941** | **3902** |
| **Clozapine, mg/kg** | **0** |  | **50** |  |

